# TSUNAMI: Translational Bioinformatics Tool Suite For Network Analysis And Mining

**DOI:** 10.1101/787507

**Authors:** Zhi Huang, Zhi Han, Tongxin Wang, Wei Shao, Shunian Xiang, Paul Salama, Maher Rizkalla, Kun Huang, Jie Zhang

## Abstract

Gene co-expression network (GCN) mining identifies gene modules with highly correlated expression profiles across samples/conditions. It helps to discover latent gene/molecular interactions, identify novel gene functions, and extract molecular features from certain disease/condition groups, thus help to identify disease biomarkers. However, there lacks an easy-to-use tool package for users to mine GCN modules that are relatively small in size with tightly connected genes that can be convenient for downstream Gene Ontology (GO) enrichment analysis, as well as modules that may share common members. To address this need, we develop a GCN mining tool package TSUNAMI (Tools SUite for Network Analysis and MIning) which incorporates our state-of-the-art lmQCM algorithm to mine GCN modules in public and user-input data (microarray, RNA-seq, or any other numerical omics data), then performs downstream GO and enrichment analysis based on the modules identified. It has several features and advantages: (i) user friendly interface and the real-time co-expression network mining through web server; (ii) direct access and search of GEO and TCGA databases as well as user-input expression matrix (microarray, RNA-seq, etc.) for GCN module mining; (iii) multiple co-expression analysis tools to choose with highly flexible of parameter selection options; (iv) identified GCN modules are summarized to eigengenes, which are convenient for user to check their correlation with other clinical traits; (v) integrated downstream Enrichr enrichment analysis and links to other GO tools; (vi) visualization of gene loci by Circos plot in any step. The web service is freely accessible through URL: http://spore.ph.iu.edu:3838/zhihuan/TSUNAMI/. Source code is available at https://github.com/huangzhii/TSUNAMI/.

## Introduction

Gene co-expression network (GCN) mining is a popular bioinformatics approach to identify densely connected gene modules, which are linked by their highly correlated expression profiles. It helps reveal latent gene/molecule interactions, identify novel gene functions, disease pathways and biomarkers, as well as provide disease mechanistic insights. GCN mining approaches such as WGCNA [1] and lmQCM [2] have been used increasingly [3–7]. Compared to the more popularly used WGCNA package, lmQCM is capable to mine smaller densely co-expressed GCN modules and allow overlapped membership in the output modules. Those features reflect closely the real biological networks *in vivo*, where the same genes may participate in multiple pathways and a small group of genes are more likely to be synergistically regulated in local pathway functions. Besides, the generally smaller size of modules from lmQCM usually generates more meaningful GO enrichment results, which has been successfully applied to many diseases and cancer types [8–17].

There exist several online databases that curate transcriptomic data, for example, PanglaoDB (https://panglaodb.se/) collected single-cell RNA-seq (scRNA-seq) data from mouse and human. Cao et al. scRNASeqDB [18] provides an scRNA-seq database for gene expression profiling in human. Recount2 [19] provides public available analysis-ready gene and exon counts datasets. However, all of these databases focus on data collection, to the best of our knowledge, there is no tool offering the entire pipeline that can directly process transcriptomic data, mine GCN modules, analyse GO enrichment, and visualized the results in a complete pipeline fashion. To meet such needs, we released our web-based analysis tool suite TSUNAMI (Tools SUite for Network Analysis and MIning).

For users’ convenience, TCGA mRNA-seq data (Illumina HiSeq RSEM genes normalized from https://gdac.broadinstitute.org/) and NCBI Gene Expression Ominbus (GEO) are directly incorporated into TSUNAMI, where GEO contains a large number of microarray datasets. In addition, other data types such as miRNA-seq and DNA methylation are also compatible with this suite. In fact, TSUNAMI can handle any numerical matrix data regardless which omics data type it is from. In TSUNAMI, it not only incorporates the newly released lmQCM algorithm, but also includes WGCNA package for users to explore and compare their GCN modules from two different algorithms. We offer highly flexible parameter choices in each step to users who may want to fine tune each algorithm to suit for their own data and goal.

Prior to data mining, a data pre-processing interface is designed to address the input data format difference and filter the data to remove noise for GCN mining. Each step of the pre-processing is transparent to users and can be adjusted according to their own preferences and needs.

Furthermore, our website directly incorporates GO enrichment analysis and Circos plot function for researchers to explore the enriched biological terms and gene locations in the output GCN modules, as well as providing a tool for survival analysis with respect to each GCN module’s eigengene values. All of the aforementioned functions only need button clicks from user-side. The design of such user-friendly implementations of our TSUNAMI pipeline provides a one-stop comprehensive analysis tool suite for biological researchers and clinicians to perform transcriptomic data analysis themselves without any prior programming skill or data mining knowledge.

## Data input

A flowchart that describes TSUNAMI pipeline is presented in **Figure 1**. The entire pipeline is implemented in R language with Shiny server pages. In the future, it will be upgraded with Python to improve the computing speed in module mining step. Some front-end interfaces and functions are done by JavaScript. In TSUNAMI, users can choose to use either TCGA RNA-seq expression data, GSE series matrix data, or other RNA-seq data from GEO database, or local user-input numerical matrix data, such as microarray, RNA-seq, scRNA-seq data, DNA methylation data, or any other type of numeric matrix data. User can also choose specific omics data type on GEO database if keywords are given to indicate the data type in the search window. Only few GSE data is not able to be processed (for example, 12 out of first 1000 GSE data), mostly are legacy microarray data, which contain too much missing data or too small sample size. Other 98.80% of first 1000 GSE data can be processed. On the website, various of example data from microarray to scRNA-seq data are listed on TSUNAMI for users’ reference. Instead of searching GEO database manually, TSUNAMI provides a friendly interface for users to retrieve data from GEO by keywords, offers flexible select tool to retrieve relevant GSE dataset to perform GCN analysis.

**Figure 1.**
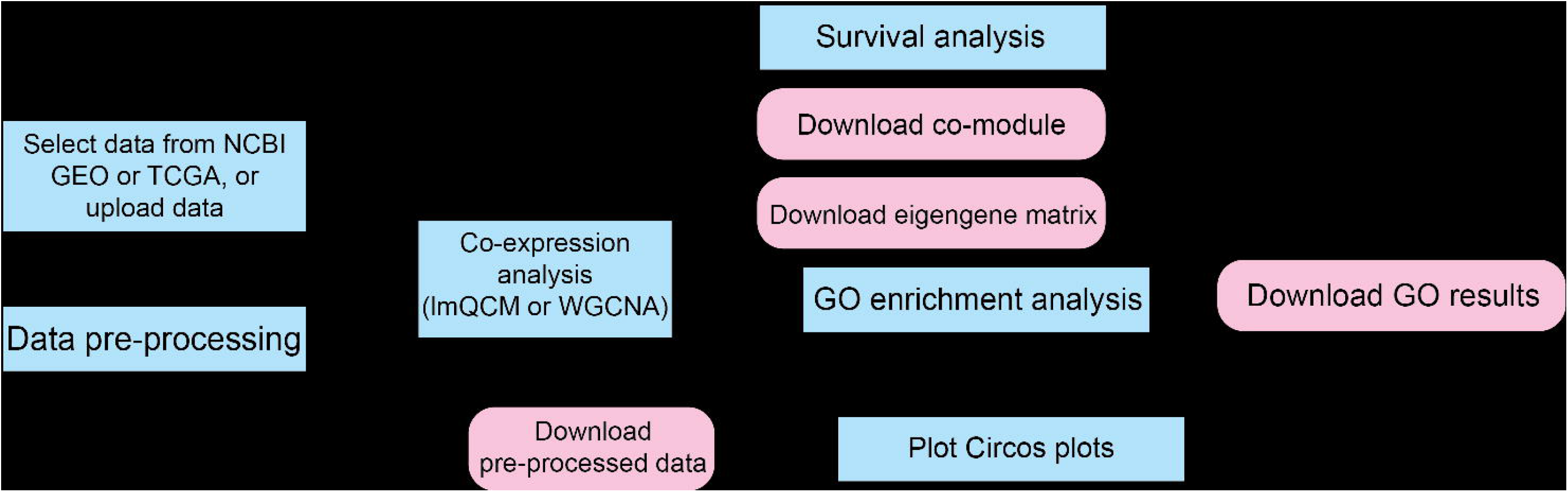
Flowchart of TSUNAMI. In this flowchart representation of TSUNAMI pipeline, blue rectangles represent pipeline operations; rounded rectangles in pink represent download processes.

TSUNAMI also provides an upload bar for users to upload local files in various formats (CSV, TSV, XLSX, TXT, etc.), the upload interface is shown in **Figure 2A**. In this paper, one microarray dataset (GSE17537 from GEO) is chosen as an example to present the features of TSUNAMI. GSE17537 contains microarray data of 55 colorectal cancer patients from Vanderbilt Medical Center (VMC), with 54,675 probesets [20, 21].

**Figure 2.**
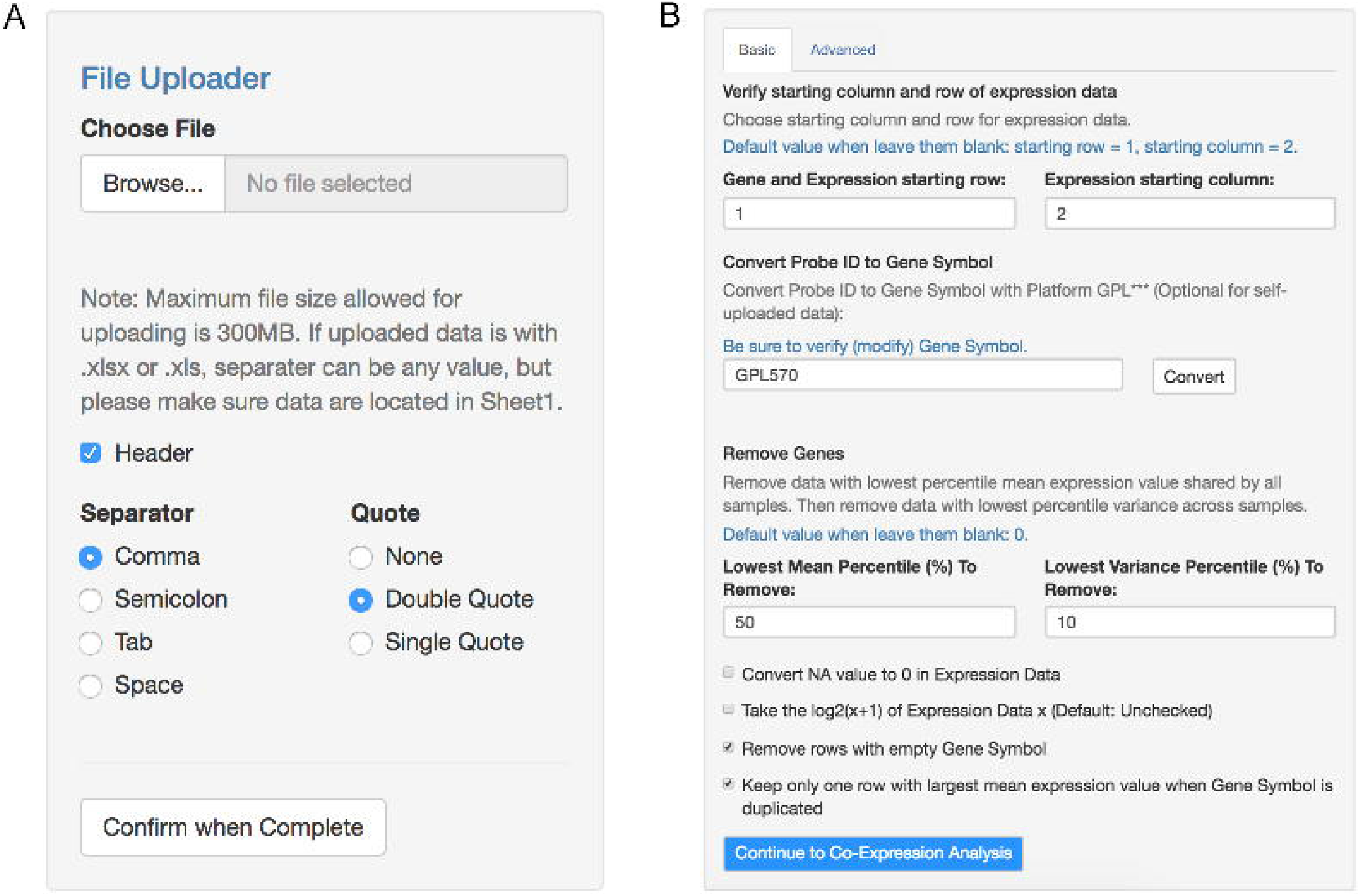
Dataset Selection and Pre-processing Panel. **A.** Data can be uploaded manually, or chosen from NCBI GEO database (not shown in the figure). When uploading the data, the maximum file size that TSUNAMI allows is 300 Megabytes. Header, separators and quote methods can be adjusted by users. **B.** The Data Pre-processing Panel includes several pre-processing steps.

### Online data pre-processing

One issue of GEO microarray data is that different platforms adopted different rules when converting probeset IDs to gene symbols. To make this step easier for users, probeset IDs in GSE data matrix from GEO can be converted to gene symbols using R package “BiocGenerics” [22] by only one click. For instance, for GSE17537, the annotation platform is GPL570. TSUNAMI can also automatically identify annotation platforms of the data from GEO. During the conversion, TSUNAMI will (i) remove rows with empty gene symbol; and (ii) select the rows with the largest mean expression value when multiple probesets are matched with the same gene symbol. The interface of data pre-processing step is shown in **Figure 2B**.

Additional data filtering steps include: (i) convert “NA” value (not a number value) to 0 in expression data, to ensure all the values are numeric and can be interpreted by co-expression algorithms; (ii) perform log_2_(*x* + 1) transformation of the expression values *x* if the original values have not been transformed previously; (iii) remove lowest *J* percentile rows (genes) with respect to mean expression values; remove lowest *K* percentile rows with respect to expression values’ variance. These data filtering steps are necessary to reduce noise and to ensure the robustness for the downstream correlational computation in lmQCM algorithm. The default settings are *J* = 50 and *K* = 50, by which genes with low expression and variance across samples are filtered out. In our example with GSE17537, we deselect logarithm conversion and NA value to 0 conversion, set *J* = 50, and *K* = 10, as shown in **Figure 2B**. However, users can always adjust these parameters based on their own needs and preferences. In Data Pre-processing section, we further provide “Advanced” panel to allow users select samples subgroup of their interest. After finished the data pre-processing, a dialog box will appear to indicate how many genes left after the filtering process.

### Weighted network co-expression analysis

After data pre-processing, users can directly download pre-processed data or further proceed to GCN analysis. In GCN analysis, we implemented lmQCM algorithm as well as WGCNA pipeline. We kept the mining steps concise and simple with default parameter settings, while preserving the flexibility for users to select parameters in each step. Guidelines for parameter selection are in method pages of the website. Besides this article, we also release the lmQCM package to CRAN (https://CRAN.R-project.org/package=lmQCM). The R package “WGCNA” from Bioconductor (http://bioconductor.org) was adopted to integrate the WGCNA pipeline.

In the lmQCM method panel, users can adjust parameters such as initial edge weight *γ*, weight threshold controlling parameters *λ*, *t*, *β*, and minimum cluster size (**Figure 3**). Pearson correlation coefficient (PCC) and Spearman’s rank correlation coefficient (SCC) are implemented separately for users to select. SCC is recommended for analysing RNA-seq data due to the large range of data values, and it is more robust than PCC to tolerate outliers. In our example with GSE17537, the default settings were used (unchecked weight normalization, *γ* = 0.7, *λ* = 1, *t* = 1, *β* = 0.4, minimum cluster size= 10, and PCC for correlation measure). The running time of lmQCM depends on the number of genes after filtering process. A progress bar is provided to show the program progress. Note that lmQCM will not work if the data contain no clustering structure or the gene pair correlations are so poor that none is above the initial mining starting threshold (*γ*). In those cases, the program will stop running and generate a warning message. However, if the data contain enough high correlated gene pairs after filtering and with the default program settings, this should not happen.

**Figure 3.**
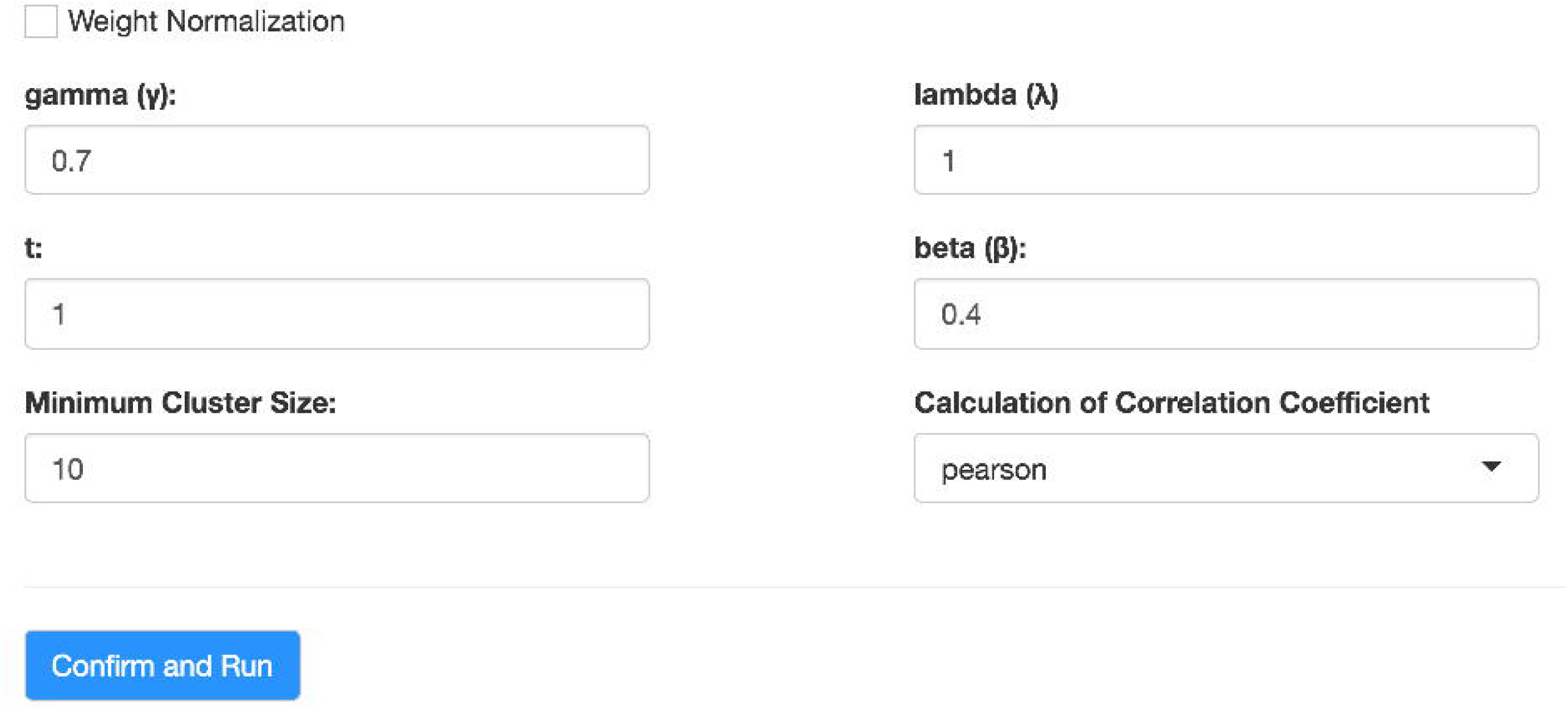
lmQCM Method Panel Data Pre-processing Panel. The lmQCM algorithm panel which allows users to choose various of parameters. In this paper, experiment runs with unchecked weight normalization, *γ* = 0.7, *λ* = 1, *t* = 1, *β* = 0.4, minimum cluster size = 10, and adopted Pearson correlation coefficient.

The WGCNA method panel is a two-step analysis: Step 1 helps users to specify the hyper-parameter “power” in step 2, *i.e.*, the soft thresholding in [1] by visualizing the resulting plot (**Figure 4A**). Step 2 allows users to select the remaining parameters. TSUNAMI allows users to customize the parameters of power, reassign threshold, merge cut height, and minimum module size. After applying WGCNA, a hierarchical clustering plot for getting the result modules is also shown in this panel (**Figure 4B**). The resulting plot in **Figure 4B** is from the example data GSE17537 with power= 10, set reassign threshold = 0, merge cut height = 0.25, and minimum module size = 10.

**Figure 4.**
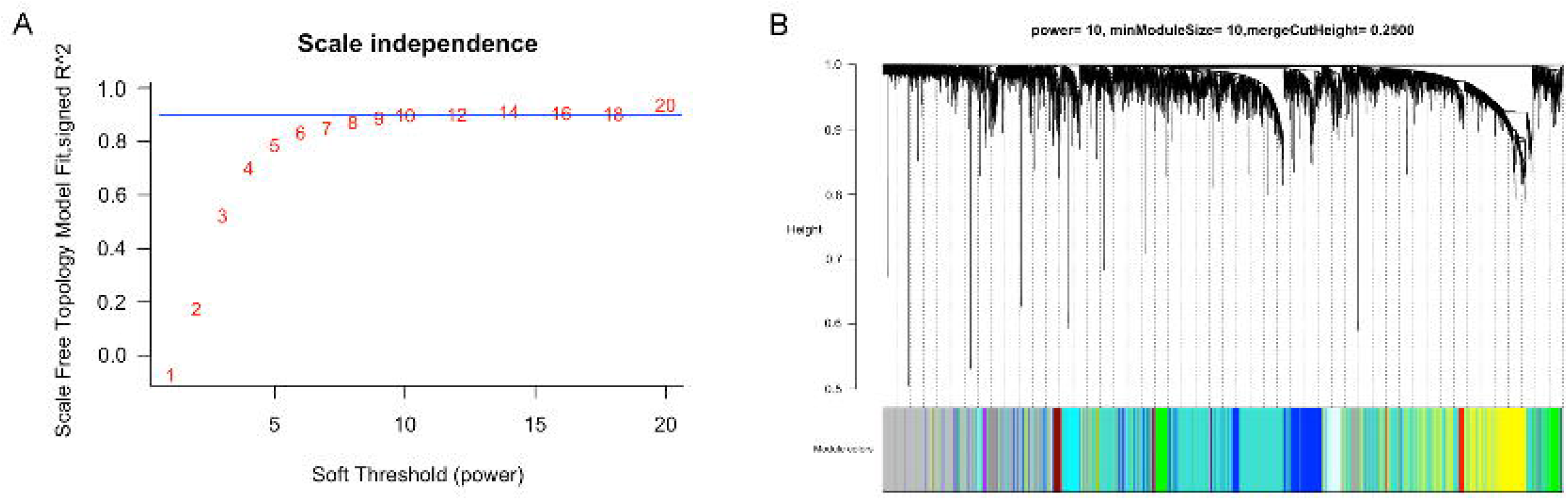
Choosing the Power in WGCNA and the Hierarchical Clustering Graph of WGCNA. **A.** The hyper-parameter “power” that chosen from the value above the blue horizontal line. **B.** The result hierarchical clustering graph with color bar indicating result modules with GSE17537 series matrix as an example, use parameters power= 10, reassign threshold = 0, merge cut height = 0.25, minimum module size = 10 in WGCNA.

In the last step of GCN mining, two outputs are provided by TSUNAMI: (i) merged gene clusters sorted by their sizes in descending order (**Figure 5A** with lmQCM algorithm); (ii) an eigengene matrix, which is the expression values of each GCN summarized into the first principal component using singular value decomposition (**Figure 5C** with lmQCM algorithm). Eigengene values can be regarded as the weighted average expressions of each GCN, thus each GCN is summarized to a “super gene” with the first right singular vector as the expression values. Such values are very useful for users to correlate GCN modules expression profiles with various traits in the downstream analysis such as survival analysis. All results can be downloaded in CSV or TXT format.

**Figure 5.**
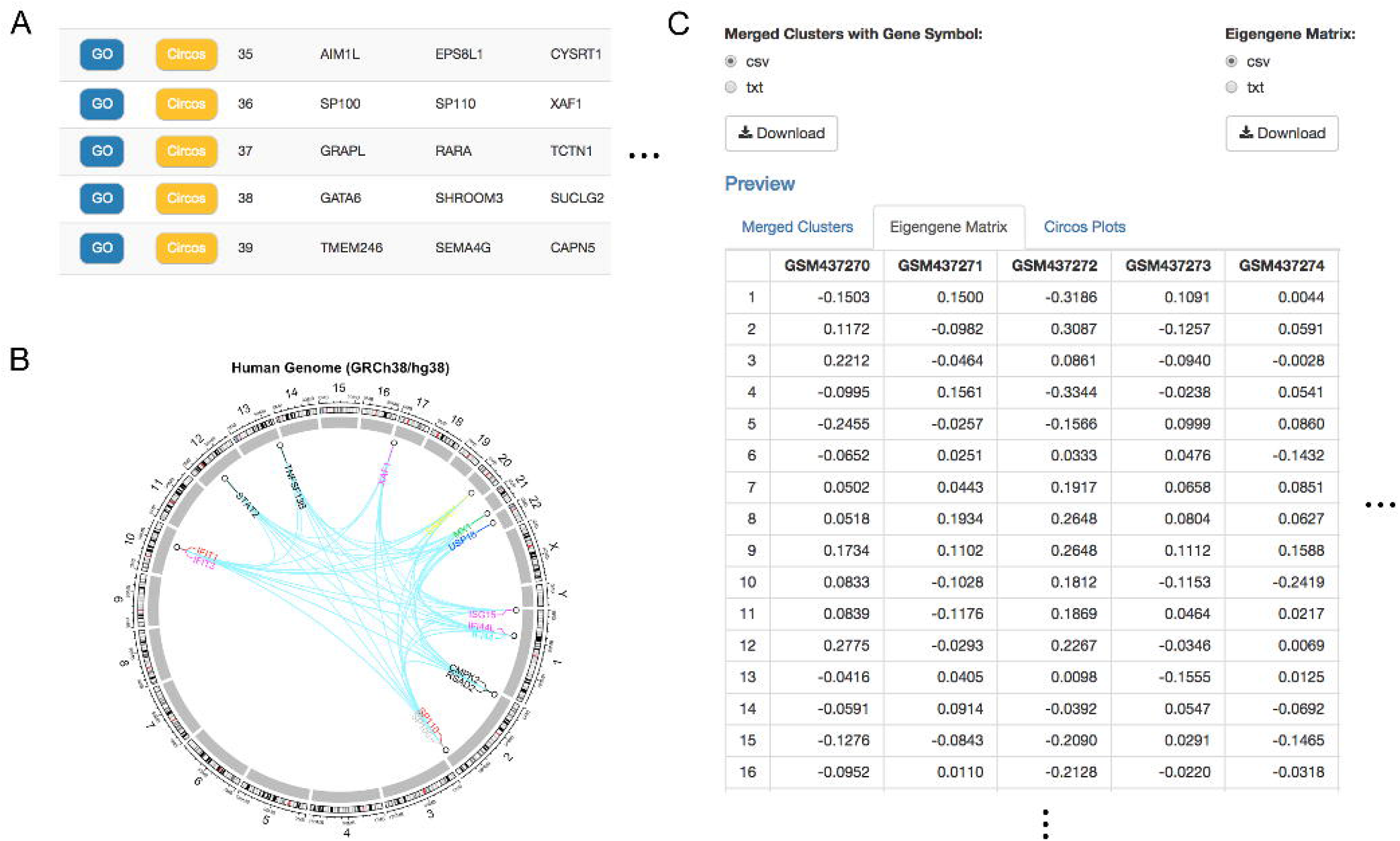
Merged Clusters Result Generated by lmQCM. **A.** The merged GCN module results, sorted in descending order based on the length of each cluster. Figure only shows part of the results (cluster 35~39) with part of genes. **B.** The Circos plot result from the 36^th^ GCN module with 15 genes. **C.** The screenshot of the eigengene matrix (rounded to 4 decimal places for better visualization). Figure only shows part of the results (cluster 1~16) with part of samples (GSM437270~GSM437274). All subfigures use lmQCM algorithm with default parameters (unchecked weight normalization, *γ* = 0.7, *λ* = 1, *t* = 1, *β* = 0.4, minimum cluster size = 10, and adopted Pearson correlation coefficient) with GSE17537 series matrix as an example.

### Downstream enrichment analysis

Enrichr [23, 24] is used as the tool for downstream GO enrichment analysis implementation. By default, total 14 types of frequent used enrichment are performed. They are (1) Biological Process; (2) Molecular Function; (3) Cellular Component; (4) Jensen DISEASES; (5) Reactome; (6) KEGG; (7) Transcription Factor PPIs; (8) Genome Browser PWMs; (9) TRANSFAC and JASPAR PWMs; (10) ENCODE TF ChIP-seq; (11) Chromosome Location (Cytoband); (12) miRTarBase; (13) TargetScan microRNA; (14) ChEA. Users can further customize the enrichment result categories from the open source code available in Github (https://github.com/huangzhii/TSUNAMI).

To access Enrichr results, users can simply click the blue button “GO” in each row adjacent to the GCN mining results (as shown in **Figure 5A**). In each enrichment analysis result, it outputs the term (e.g., GO or pathway), *P* value, z-score, overlapped genes, etc. Users can download multiple analysis results which are bundled in a ZIP file. Besides, other popular GO analysis websites are also directly linked in TSUNAMI to bring conveniences to users. In our example with GSE17537, we select the 36^th^ GCN module with 15 genes generated by lmQCM to analyze the GO enrichment, and each result table are sorted based on the *P* value that Enrichr calculated. From the result in **Table 1**, we can see the 36^th^ GCN module is highly overlapped with GO Biological Process term “type I interferon signaling pathway (GO:0060337)” (9 out of 148 genes).

### Circos plot

TSUNAMI provides Circos plots [25] through any intermediate results or inputs in the cases of human transcriptomic data. Circos plot is a very useful graph for visualizing the positions of genes on chromosomes and gene-gene relationships/interactions. The Circos plot function from the R package “circlize” [25] is adopted in this package for users to locate and visualize mined GCNs of human genes.

In TSUNAMI, users can visualize the Circos plot via “Circos Plots” section, either by typing their own genes list separated by carriage return character (“\n”) directly, or using the calculated GCN modules (for example, by clicking the yellow button right next to the “GO” button in **Figure 5A**). TSUNAMI supports both human genomes hg38 (GRCh38) and hg19 (GRCh37). To match the gene symbol to chromosomes’ starting and ending sites, we use reference gene table from UCSC genome browser [26]. If multiple starting/ending site are matched, we choose the longest one with length calculated by:

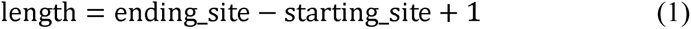

By updating the plots, users can also choose the size of the plots and decide whether gene symbols and pair-wised links should be shown on the graph.

An example output of Circos plot in **Figure 5B** used the 36^th^ GCN module with 15 genes in the lmQCM result from GSE17537 series matrix (use a color set for texts to get a clear visual effect), indicated by gene symbols of human genome hg38 (GRCh38). While the link between a pair of genes indicates that they belong to the same co-expressed GCN module.

Circos plots can help users to visualize the GCN module’s location on human chromosomes from either lmQCM or WGCNA mining, help them to visualize GCNs due to copy number variation and other structural changes. In the future, genome from mouse and other species will be incorporated for Circos plot.

### Survival analysis with respect to GCN modules

An optional step of survival analysis follows the generation of the eigengene matrix. It allows users to correlate the GCN module’s eigengene values with patient clinical survival (or event-free survival), and such extension tool can be further customized as users’ need to correlate module eigengene values with other clinical traits in the future version. In our current version, we only implemented survival analysis as an example.

In the survival analysis, users can perform Overall Survival/Event-Free Survival (OS/EFS) analysis based on the GCN modules’ eigengene values, and look for significant GCNs that are capable for prognosis, although depending on the group of patients user specifies, such GCNs may not be identified all the time. TSUNAMI lets user to select an eigengene row (corresponding to a GCN module). The program will splits the patients into two groups by eigengene values’ median, then tests two groups against OS/EFS by calculating the *P* value of the log-rank test [27, 28]. Before doing so, users need to input the numerical survival time of OS/EFS (either in months or in days) with categorical events OS/EFS status (1: deceased; 0: censored). “survdiff” function from R package “survival” is adopted to calculate the *P* value and plot the Kaplan-Meier survival curve.

Take GSE17537 with full survival information as an example, the Kaplan-Meier survival plot is generated according to the OS information by dichotomizing the 36^th^ GCN module’s eigengene values by its median to high and low group, as shown in **Figure 6**. Such GCN module was generated from lmQCM method with default settings as shown in **Figure 3**. This survival analysis offers researchers the tool to immediately identify any GCN modules that reflects patients’ survival difference, thus allows researchers to further study their roles as potential prognosis biomarkers, as well as the biological pathways that differentiate the patients.

**Figure 6.**
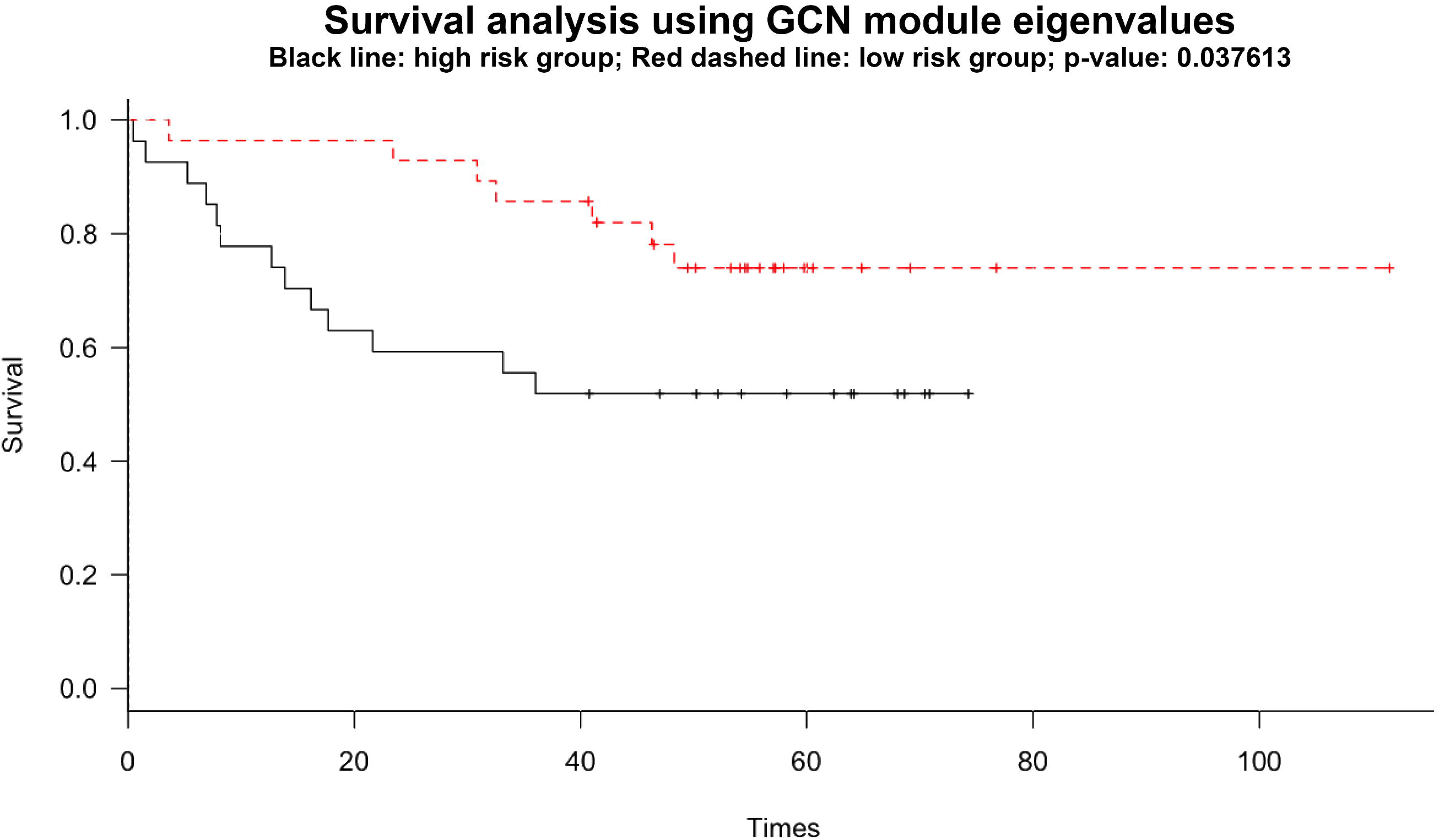
Survival Analysis using GCN Module Eigenvalues. Survival analysis using the 36^th^ GCN module eigenvalues generated from lmQCM algorithm, with default parameters (unchecked weight normalization, *γ* = 0.7, *λ* = 1, *t* = 1, *β* = 0.4, minimum cluster size = 10, and adopted Pearson correlation coefficient) with GSE17537 series matrix as an example. 55 samples are used with Overall Survival information.

## Conclusion

We released the TSUNAMI online tool package for gene co-expression modules identification with direct link to TCGA RNA-seq database and GEO transcriptomic database as well as users’ input data. It is a one-stop comprehensive tool package which has several advantages such as flexibility of parameter selections, comprehensive GCN mining tools, direct link to downstream GO enrichment analysis, Circos plot visualization, and survival analysis, with downloadable results in each step. All of which bring tremendous convenience to biological researchers.

Besides, TSUNAMI can not only process microarray, RNA-seq, and single-cell RNA-seq transcriptomic data, but also be capable for processing any type of the numerical valued matrix for weighted network module mining. If the users upload an adjacency matrix of any supported format with numerical values as the edge weights, TSUNAMI can be used to mine any correlational network modules or even beyond that. This extension will be implemented in version 2.0.

**Table 1 The partial results of GO enrichment analysis**

*Note*: This table contains partial rows and columns from original result (active panel: GO Biological Process) from the 36^th^ GCN module with 15 genes generated by lmQCM with GSE17537 series matrix as data. GO terms are sorted by *P* value. We refer readers to explore other *P* values and scores from TSUNAMI webpage and Enrichr package.

## Authors’ contributions

JZ and KH conceived the idea of the project and participated in software design and helped to draft the manuscript. Zhi Huang and Zhi Han wrote the software and manuscript. TW, WS, and SX carried out the GO enrichment analysis tool options. PS, MR, KH, JZ provide research guidance. JZ and KH reviewed and edited the manuscript. All authors read and approved the final manuscript.

## Acknowledgements

This work is partially supported by IUSM startup fund (JZ), ACS IRG grant (JZ), the NCI ITCR U01 (CA188547) grant (JZ and KH), and Indiana University Precision Health Initiative (JZ and KH). We thank the support from Indiana University Information Technologies and Advanced Biomedical IT Core. The results shown here are partially based upon data generated by the TCGA Research Network: https://www.cancer.gov/tcga.

