## Supplementary material for "TSUNAMI: Translational Bioinformatics Tool Suite For Network Analysis And Mining": Table 1

**Table 1 The partial results of GO enrichment analysis**

| **ID** | **Term** | **Overlap** | ***P* value** | **Z-score** | **Genes** |
| --- | --- | --- | --- | --- | --- |
| 1 | Type I interferon signaling pathway (GO:0060337) | 9/148 | 2.51E-16 | -3.2821 | SP100; RSAD2; STAT2; MX1; ISG15; SAMHD1; XAF1; IFIT1; IFIT3 |
| 2 | Cellular response to type I interferon (GO:0071357) | 4/23 | 1.80E-09 | -2.7766 | SP100; MX1; ISG15; IFIT1 |
| 3 | Negative regulation of single stranded viral RNA replication via double stranded DNA intermediate (GO:0045869) | 4/44 | 2.73E-08 | -2.6829 | RSAD2; MX1; ISG15; IFIT1 |
| 4 | Negative regulation of viral genome replication (GO:0045071) | 4/40 | 1.84E-08 | -2.4940 | RSAD2; MX1; ISG15; IFIT1 |
| 5 | Negative regulation by host of viral genome replication (GO:0044828) | 4/51 | 5.01E-08 | -2.6224 | RSAD2; MX1; ISG15; IFIT1 |
| 6 | Response to type I interferon (GO:0034340) | 3/35 | 2.20E-06 | -2.7155 | SP100; MX1; ISG15 |
| 7 | Regulation of type I interferon-mediated signaling pathway (GO:0060338) | 3/43 | 4.14E-06 | -2.6859 | STAT2; SAMHD1; USP18 |
| 8 | Negative regulation of type I interferon-mediated signaling pathway (GO:0060339) | 2/43 | 4.66E-04 | -2.5488 | STAT2; USP18 |
| 9 | Positive regulation of type I interferon-mediated signaling pathway (GO:0060340) | 2/52 | 6.81E-04 | -2.5122 | STAT2; USP18 |
| 10 | Positive regulation of Fas signaling pathway (GO:1902046) | 1/7 | 5.24E-03 | -2.9563 | SP100 |

*Note:* This table contains partial rows and columns from original result (active panel: GO Biological Process) from the 36^th^ GCN module with 15 genes generated by lmQCM with GSE17537 series matrix as data. GO terms are sorted by *P* value. We refer readers to explore other *P* values and scores from TSUNAMI webpage and Enrichr package.
